# Day length-dependent thermal COP1 dynamics integrate conflicting seasonal cues in the control of *Arabidopsis* elongation

**DOI:** 10.1101/2021.11.25.470009

**Authors:** Cristina Nieto, Pablo Catalán, Luis Miguel Luengo, Martina Legris, Vadir López-Salmerón, Jean Michel Davière, Jorge Casal, Saúl Ares, Salomé Prat

**Author notes:** These two authors contributed equally to this work (Cristina Nieto, Pablo Catalán).

## Abstract

As the summer approaches, plants experience enhanced light inputs and elevated temperatures, two environmental cues with an opposite morphogenic impact. How plants integrate this conflicting information throughout seasons remains unclear. Key components of the plant response to light and temperature include phytochrome B (phyB), PHYTOCROME INTERACTING FACTOR 4 (PIF4), EARLY FLOWERING 3 (ELF3) and CONSTITUTIVE PHOTOMORPHOGENIC 1 (COP1). Here, we used hypocotyl lengths of single and double mutant/over-expression lines to fit a mathematical model incorporating known interactions of these genes. The fitted model recapitulates day length-dependent thermoelongation of all lines studied, and correctly predicts temperature responsiveness of new genotypes. Whilst previous works pointed to a light-independent thermal function of COP1, simulations of our model suggested that COP1 has a role in temperature-signaling only during daytime. Based on by this prediction, we show that COP1 overexpression increases thermal response in continuous white light, while it has little effect in darkness. Defective thermal response of *cop1-4* mutants is epistatic to *phyB-9* and *elf3-8*, indicating that COP1 activity is essential to the transduction of phyB and ELF3 thermosensory function. Our model accurately captures phyB, ELF3 and PIF4 dynamics, providing an excellent toolbox for identification of best allelic combinations towards optimized crops resilience to climate change at different geographical latitudes.

## INTRODUCTION

Light and temperature are key environmental factors that shape plant growth patterns according to the prevailing conditions of the environment. Growth of the hypocotyl, for instance, proceeds very rapidly in young *Arabidopsis* seedlings buried in the soil, to speed up exposure of photosynthetic organs to sunlight. Upon emergence, light severely reduces the hypocotyl growth rate, while this is reassumed on seedlings exposure to neighboring vegetation that threat resources availability. Warm temperatures alike enhance hypocotyl growth to facilitate cooling of aerial tissues. The choice of sowing date is a critical decision in crop management to optimize overlap with the most favorable season. As sowing dates progress towards the warmer season, plants sense the increased light input of longer day lengths. However, how contrasting effects of light and temperature integrate to the control of thermomorphogenic development across seasons remains poorly understood.

The predominant photoreceptors inhibiting hypocotyl growth in *Arabidopsis* are phytochrome B (phyB), phyA, and cryptochromes. phyB acts as a main thermosensor, featuring a cross-talk interaction of light and temperature information already at the sensor level. phyB is synthetized in an inactive form (Pr) and photo-converted upon red-light (R) perception into the active Pfr state that is translocated into the nucleus for light signaling transduction ^1^. Far red-light (FR) rapidly reverses Pfr into the Pr state and through this conversion phyB senses changes in the R/FR ratio that result from neighboring plants ^2,3^. Moreover, phyB Pfr slowly returns into the inactive Pr form, in a process termed thermal or dark reversion that is accelerated by warm temperatures ^4,5^. While phyA and cryptochromes have a main role in light-induced transcriptional reprogramming, so far there is no evidence for these photoreceptors acting as temperature sensors.

ELF3 confers the circadian clock’s “evening complex” (EC) thermal responsiveness via a prion-like thermosensor domain that reversibly directs protein phase transition ^6^. ELF3, ELF4 and LUX occupancy of their target loci is strongly reduced at warmer temperatures^7,8^, with loss of *elf3, elf4*, or *lux* function leading to increased growth at 22°C, indicative of a constitutively activated thermal response in these mutants ^9^. Temperature-induced elongation is arbitrated by transcriptional activation of cell wall loosening and auxin biosynthetic/ signaling genes by the PIF4 and PIF7 factors ^10,11^. Translation of PIF7 is increased at warm temperatures through structural changes of an RNA hairpin loop within the 5’ untranslated transcript region, with this control likely having a prevalent role on daytime activation of the thermomorphogenic pathway under longer day lengths.

Light activated phyB suppresses PIF4 activity by signaling its phosphorylation and subsequent degradation by the 26S proteasome, in addition to prevent PIF4 binding to its target promoters ^12,13^. The circadian clock regulates rhythmic *PIF4* expression via transcriptional repression by the LUX, ELF4 and ELF3 proteins, comprising the core evening complex loop ^14^. The EC is recruited by the LUX factor to the *PIF4* promoter, suppressing *PIF4* expression during early night ^15^.

Additionally, phyB Pfr inactivates the E3 ligase COP1, which acts as a photomorphogenesis suppressor in the dark. COP1 localizes into the nucleus in darkness, where it targets degradation of multiple photomorphogenesis-promoting factors, including LONG HYPOCOTYL IN FAR-RED 1 (HFR1) and ELONGATED HYPOCOTYL (HY5) ^16–18^. Red-light dependent phyB and SPA1 interaction dissociates the SPA1-COP1 complex and suppresses COP1 activity ^19^, besides triggering COP1 nuclear exclusion through a yet elusive mechanism ^20^. HFR1 and HY5 promote photomorphogenesis by antagonizing PIF4, through formation of inactive HFR1-PIF4 complexes and competitive interaction of HY5 with the PIF4 cognate elements ^21–24^. Elevated temperatures promote in turn COP1 nuclear accumulation ^25^, hence alleviating HY5 suppressive effects. COP1 is also reported to trigger ELF3 destabilization during ELF3-dependent recruitment of GIGANTEA (GI) for COP1 degradation ^26^, although relevance of this process in growth responses remains unknown.

Together, these observations reveal that phyB, ELF3, PIF4 and COP1 form a thermal network where all of these regulators physically and functionally interact with each other. Along with phyB suppression of PIFs and COP1 activity, COP1 is shown to facilitate phyB Pfr turnover under extended light ^27^. Furthermore, phyB favors ELF3 accumulation presumably by suppressing protein destabilization by COP1 ^28^, besides co-occupying at cooler temperatures many of the EC target loci ^7^. PIFs, in turn, promote COP1-mediated phyB degradation ^27^, and recruit phyB into the Light-Response BTB1 and 2 (LRB1 and LRB2) complex, involved at phyB and PIFs ubiquitination and mutual destruction ^29,30^ Thus, extensive connectivity of these regulators makes them prime candidates for modeling studies, to dissect their exact roles in thermal elongation and assess whether their thermosensory function is dependent on light.

Here we investigated the output of this network in seedlings grown under different day lengths and 22°C/28°C, as a mimic of seasonal information. We used the hypocotyl lengths of mutants and overexpresser lines to fit a mathematical model based on the molecular interactions of these regulators. We validate the predicted molecular dynamics of these components and use the generated model to weigh their relative contribution to thermal elongation as influenced by day length.

## RESULTS

### Day length-dependent roles of phyB, ELF3, and COP1 in thermal elongation

To gain a better understanding on the combined roles of phyB, ELF3, and COP1 in coordinating thermomorphogenesis (Fig.1A), we examined temperature-induced elongation of single and double mutants and overexpression lines in the Col-0 background. Light effects on thermal responsiveness were analyzed by measuring their hypocotyl lengths under continuous dark, continuous white light (CWL), and 8h, 12h and 16h light cycles, at a temperature of 22°C or 28°C (Fig.1A, Material and Methods).

**Figure 1.**
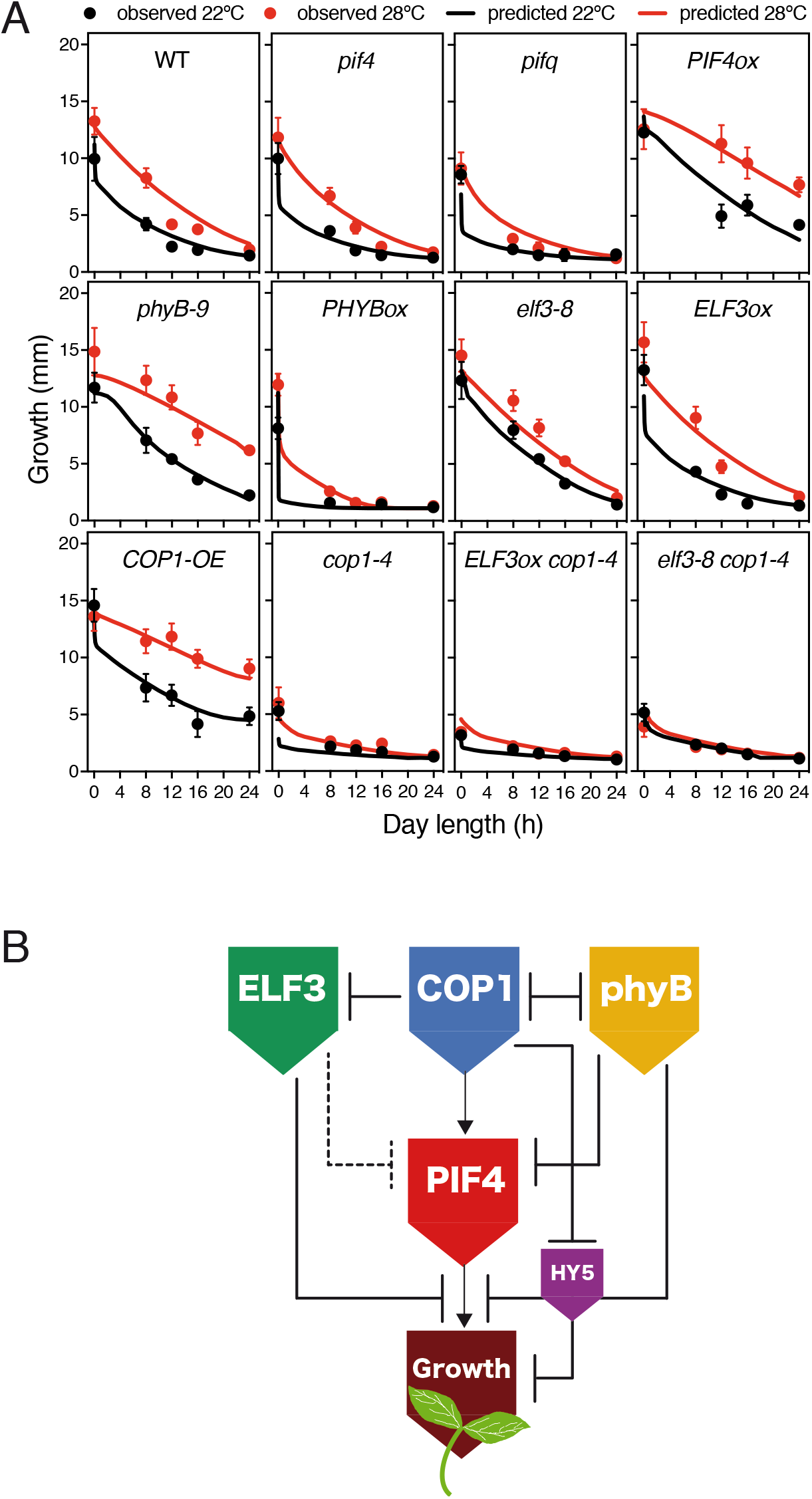
Phenotype of loss-of function mutants and overexpressors under different temperatures and day lengths. **(A)** Growth of twelve *Arabidopsis* genotypes at either 22°C or 28°C under six white light day length conditions. Measured hypocotyl lengths (circles) are compared with values predicted by the growth model (solid lines). Bars indicate standard deviation (n=30). The model accurately captures the main trends of data. **(B)** Schematic representation of the network tested. Dashed and solid lines indicate transcriptional and post transcriptional regulations respectively.

Thermal response of wild-type plants was found in these studies to be inversely correlated with day length. Hypocotyl elongation was greatest at 8 h light and progressively reduced by 12 h and 16 h day lengths, while Col-0 seedlings lacked any thermal response in CWL (Fig.1A). Thermal elongation was also significantly reduced in continuous darkness, suggesting that thermal growth requires of initial phyB Pfr photoconversion and photomorphogenic induction, in addition to alternating light/dark cycles. This is consistent with gating by the circadian clock to be pivotal to this response. Thermal growth was moreover suppressed in *pif4* and *pifq* mutants (Fig.1A), while it was notably enhanced in *phyB-9* mutant and *PIF4ox* lines, in agreement with phyB acting at temperature perception ^4^ and a downstream role of PIF4 in promoting thermal elongation ^10^. However, as opposed to the inhibitory effects of increasing light hours in the wild-type and *pif* mutants, longer day lengths enhanced thermal responsiveness of the *phyB-9* and *PIF4*ox lines. These plants showed almost maximal thermoelongation in CWL, consistently with temperature inactivating phyB Pfr also during daytime ^5^.

Thermal response was impaired in all *cop1-4* genotypes, demonstrating that COP1 activity is essential to thermal responsiveness. These plants are much shorter in darkness than *pif4* and *pifq* mutants, in line with the strong constitutive photomorphogenic phenotype of this weak allele. Thermal sensitivity defects were also more severe than for *pif4* and *pifq* mutants, with *cop1-4* seedlings failing to elongate at warmer temperatures, whereas *pif4* mutants elongated in shorter day lengths. More importantly, COP1 over-expression caused a similarly enhanced thermoresponse in CWL as seen in *phyB-9* mutants and *PIF4ox* lines, indicating that COP1 turnover of PIFs-antagonizing factors like HY5 is pivotal to thermomorphogenic growth. A greater thermal response in CWL was also observed in the *hy5* mutant (Supp. Fig. 1A), consistent with targeted HY5 turnover being to a large extent responsible of this phenotype ^31^. *COP1-OE* lines were also more elongated in CWL than the *phyB-9* mutant, suggesting that COP1-dependent PIFs stabilization ^32,33^ is biologically more relevant in prolonged light. Alternatively, other phytochromes may suppress *phyB-9* elongation.

Although PIF4 was reported to play a prominent role in driving thermomorphogenesis, *pif4* mutants showed a wild-type thermal response in darkness or shorter day lengths (Tukey’s HSD test, Fig.1A), presumably due to temperature responsive PIF7 activity ^11^. Thermal sensitivity was however decreased in *pifq* mutants, indicating that PIF1, PIF3 and PIF5 contribute as well to thermal growth. Hypersensitive response to day length of *PHYB*ox lines is indeed associated with suppressed thermomorphogenesis, indicating that thermal phyB Pfr inactivation leads to stabilization of all PIFs. Furthermore, *PHYBox* lines show remarkable thermoelongation in darkness, underscoring a light independent function of other thermosensors.

As earlier reported, *elf3-8* mutants showed de-repressed thermal elongation at 22°C ^9^, though a residual response was still observed in all tested conditions (Tukey’s HSD test, Fig.1A). Sensitivity to day length was the most significant among all assayed genotypes but it did not change with temperature, hence indicating that gating effects of the clock are important to mediate suppression of thermal responsiveness in longer day lengths. Unlike *PIF4ox* lines, growth of *elf3-8* mutants was inhibited in CWL, suggesting that ELF3 has other functions than gating *PIF4* expression ^15^. In this regard, *ELF3ox* lines were impaired in thermal elongation in long days (16 h light), though they behaved as the wild type in other diel conditions. These plants were as tall in darkness as *elf3-8* mutants, showing that interaction with phyB Pfr is key to ELF3 function ^34^.

*ELF3ox cop1-4 and elf3-8 cop1-4* lines exhibited an identical photomorphogenic development and defective thermal elongation as *cop1-4* mutants. *ELF3ox cop1-4* lines were somehow shorter than *cop1-4* in darkness, in contrast to the tall hypocotyls of etiolated *ELF3ox* (Fig.1A). This phenotype suggests that COP1 antagonizes ELF3 presumably by targeting degradation of the protein, whereas light may partially suppress ELF3 turnover.

Together, these data reveal that phyB, COP1 and ELF3 govern thermal elongation under diel conditions in a highly intertwined fashion (Fig. 1B). Dynamic changes in phyB Pfr levels as a result of external light and temperature cues drive thermal growth by suppressing PIFs nuclear accumulation, along with COP1-mediated turnover of HY5, and presumably ELF3. On the other side, thermal elongation of *phyB-9* and *PHYB*ox lines in the dark reveals that ELF3 and possibly COP1 have a role in modulating thermal responsiveness, independently of phyB Pfr.

### A mathematical model on activity of these regulators captures thermal growth of all genotypes

To assess biological significance of these regulatory events, measured hypocotyl lengths were used to fit the parameters of a mathematical model built on the known interactions of these regulators (Fig. 1B).

We used four differential equations (Supplementary Material, Eq. (S1)) that captured active levels of phyB, PIFs, ELF3 and COP1, and a fifth one that represented hypocotyl growth. For phyB dynamics we adhered to the thermal reversion model previously described ^4^, where phyB is photoactivated in the light but spontaneously reversed to its inactive Pr form in the dark, whereas the rate of phyB inactivation is temperature dependent. Dynamics of ELF3 was modelled based on its circadian control, with *ELF3* transcripts following a broad peak of expression at dusk ^15,35^. COP1 was modeled in quite an agnostic way as its nuclear levels depend on interactions with several photoreceptors for which we had no experimental information ^36^.

PIFs activity was modelled according to ELF3 repressing *PIF4* and *PIF5* transcription, with two additional terms incorporating phyB inhibition of PIFs activity, and COP1-dependent stabilization of the PIF proteins ^30,32^. Finally, hypocotyl growth was represented as the PIFs-dependent activation of growth-related genes^37^, and the competitive effects of ELF3, phyB, and HY5, on PIFs occupancy/transcriptional activation of these genes ^1^. No additional equation was included for HY5, to which its steady state levels were assumed to mainly depend on proteolytic degradation by COP1 ^39^. A full mathematical definition of the model is provided in Supplementary Material.

A custom simulated annealing algorithm (Supplementary Material) was used to fit these different equations to the growth dataset generated for the various genotypes (Methods). The fit showed that several parameters of the initial full model, Eq. (S3), were consistent with values of zero, so we proceeded to simplify the equations by eliminating these terms, Eq. (S1). In particular, the model worked equally well excluding the effect of COP1 on phyB degradation and PIFs’ action on phyB desensitization ^27^. This does not mean that these interactions do not exist, but rather that they are quantitatively less relevant than others in our experimental conditions, or that their effect is captured by other terms. The final model reproduced the dominant trends of all genetic backgrounds (Fig. 1A, solid lines), demonstrating that dynamic control of these regulators is sufficient to explain their thermomorphogenic phenotypes. Moreover, it allowed extrapolating their thermal behavior in other day length conditions (Supp. Fig. 1A), in addition to correctly predict thermal responsiveness of new genotypes (Supp. Fig. 1B). The model predicts, for instance, that suppressed thermal responsiveness of *PHYB*ox lines is partly restored under 4h light. Validation of this behavior (Supp. Fig. 1B) therefore shows that thermal Pfr inactivation reverses growth inhibition by excessive Pfr conversion in the light. The model also estimates that thermal responsiveness of double *phyB-9 elf3-8* mutants linearly increases with day length, as seen in *phyB-9*. Experimental verification of this phenotype supports an additive effect of ELF3 and phyB on thermomorphogenesis (Supp. Fig. 1A). We also validated the *cop1-4* alike phenotype of *phyB-9 cop1-4* mutants, showing that enhanced COP1 action driven by Pfr thermal inactivation is key to thermomorphogenic growth (Supp. Fig. 1A). All code and data used for fitting and model simulations can be found at https://github.com/pablocatalan/hypocotyl.

### Temperature-dependent dynamics of ELF3 and PIF4

ELF3 is a key node required for transmitting temperature information to the regulated gating of *PIF4* and *PIF5* expression at night ^40^. We therefore used the luciferase reporter p*ELF3*::LUC, p*ELF3*::ELF3-LUC, p*PIF4*::LUC and p*PIF4*::PIF4-LUC lines for the non-invasive imaging of temperature effects on *ELF3 and PIF4* peak expression and protein levels. Lines expressing the ELF3-LUC and PIF4-LUC fusions showed identical protein profiles as p*ELF3*::ELF3-myc and p*PIF4*::PIF4-HA lines (Supp. Fig. 2A and 2F), thereby allowing the analysis of rhythmic changes in these proteins during several consecutive days (Fig. 2A and 2B; Methods).

**Figure 2.**
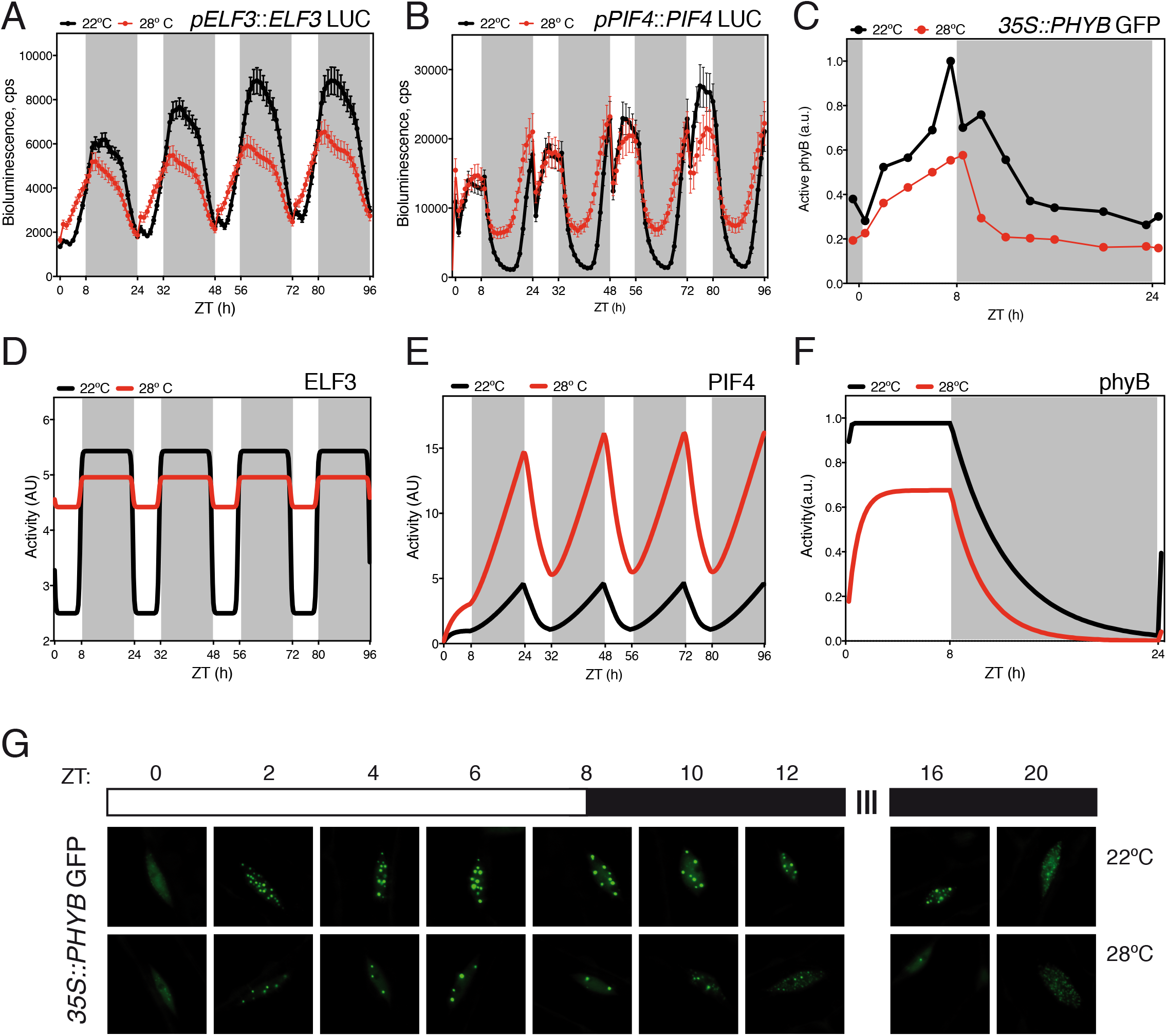
Predicted and observed dynamics of ELF3, PIF4 and phyB. Bioluminescence assays of Col-0 lines expressing constructs p*ELF3::*ELF3-LUC **(A)** and p*PIF4::*PIF4-LUC **(B)**. Seedlings were grown in short day cycles at the indicated temperatures. Values represent mean ± SEM of absolute bioluminescence of at least 24 seedlings. Bioluminescence of seedlings was measured every hour. **(D)** ELF3 and **(E)** PIF4 activity predicted by the model. **(C and G)** Temperature effects on phyB nuclear bodies in short day conditions. phyB-GFP transgenic seedlings were grown in short day cycles at either 22°C or 28°C for 5 days. Mean size of phyB nuclear bodies and total nuclear fluorescence measured with the Matlab software from confocal images **(C)** and diagram of sampling time points and growth conditions **(G)**. Arbitrary units (a.u.) of phyB activity were calculated by multiplying the nuclear bodies mean size by nuclear fluorescence at each sampling time point. Ten seedlings were measured in each assay and two biological replicates were analyzed. Warmer temperature results in fewer photobodies and less nuclear intensity. All seedlings were grown under 50 μmol.m ^-2^·s^-1^ white light in short day cycles at 22°C or 28°C. **(F)** phyB activity predicted by the model. Rectangles indicate light conditions: white, lights on, and grey, lights off. ZT indicates the zeitgeber time.

Consistent with previous reports ^15,35^, p*ELF3*::LUC expression showed a robust circadian rhythm peaking during nighttime (Supp. Fig. 2B and 2D). A roughly equivalent pattern was observed for the ELF3-LUC protein, except by showing some accumulation in short days in the light (Fig. 2A). Warm temperatures dampened p*ELF3* activity and slightly advanced its oscillation phase. Moreover, p*ELF3* waveform switched in short days to a steady state pattern, whereby LUC activity attained a constant plateau on dark transition and returned to basal levels by end of night (Supp. Fig. 2B). Interestingly, reduction of ELF3-LUC activity was lesser than for p*ELF3* transcription, indicating that ELF3 accumulates at warmer temperatures (Fig. 2A). ELF3-LUC bioluminescence was higher at 28°C during the day, and did not decline on dark transition as observed at 22°C. The same trend is seen in western blots conducted with p*ELF3*::ELF3-myc lines (Supp. Fig. 2A), implying that temperature inactivates the EC via other molecular mechanisms than the direct control of ELF3 protein stability.

In short days, *pPIF4* expression increased before dawn, as earlier described ^15^. Its transcription peaked during daytime and rapidly declined at night (Supp. Fig. 2G). Consistently with temperature suppressing gating by the EC ^40^, LUC levels were elevated at 28°C earlier at night (Supp. Fig. 2G). In line with previous reports ^41^, *pPIF4* was however expressed in long days exclusively during daytime (Supp. Fig. 2I). Also, temperature did not lead to precocious p*PIF4* up-regulation as in short days (Supp. Fig. 2I), but enhanced the amplitude and delayed *pPIF4* phase. This suggests that the EC gates *PIF4* expression during long day afternoon, as opposed to its nighttime repressive action in short days (Supp. Fig. 2G and 2J).

PIF4-LUC levels had a similar profile as *pPIF4* expression. The protein was in short days stabilized by end of the night (Fig. 2B), while in long days it decayed earlier than p*PIF4* expression during late afternoon (Supp. Fig. 2H). Also, premature decay in PIF4-LUC levels is lost at 28°C, where PIF4-LUC and *pPIF4* activities essentially overlap (Fig. 2E). Thus, PIF4 protein stabilization during long days late afternoon/ early night may provide a window for thermal elongation, in the absence of elevated *PIF4* expression at night.

Remarkably, although we implemented ELF3 activity as oscillating between two stable states (Supplementary Material), the model captures the ELF3 protein dynamics we observed experimentally (Fig. 2A, 2D and Supp. Fig. 2E). Total ELF3-LUC levels were greater in the day at 28°C, whereas differences in protein levels diminished during nighttime, consistent with the model’s prediction. Likewise, the model predicts PIF4 activity to be greater at night, and significantly enhanced at 28°C (Fig. 2E). PIF4-LUC abundance in fact increases at 28°C during nighttime or late afternoon (Fig. 2B and Supp. Fig. 2F), although it is also elevated in the day as seen in previous reports ^41–43^. This PIF4 pool is admitted to be mostly inactive due to inhibition by phyB, HY5, or the DELLA repressors (Park *et al*., 2017; Johansson *et al*., 2014; de Lucas *et al*., 2008), the model thereby correctly estimating active PIF4 levels.

### Diurnal control of phyB thermal reversion

Reduced steady-state phyB Pfr levels at warmer temperatures lead to the disassembly of large phyB photobodies, thought to be active sites for phyB signaling ^5,45^. Formation of these subnuclear domains in bright light tightly correlates with inhibition of hypocotyl growth ^46^, whereas increased Pr/Pfr ratios, in the shade or dim light, give rise to phyB redistribution into many small foci and the nucleoplasm, thereby providing an excellent cell biology readout for phyB activity ^47^. We used *35S*::PHYB-GFP lines to monitor day length and temperature contribution to phyB nuclear dynamics. Plants were grown in short days and 22°C or 28°C for 4 days and, on the fifth day, confocal images were obtained from the middle hypocotyl every two hours, for a 24 h interval (Fig. 2G). The number and size of nuclear bodies were quantified in these images (Supplemental Fig. 3, Methods), and active phyB levels (Fig. 2C) extrapolated by multiplying total nuclear fluorescence (Supp. Fig. 3A) by nuclear bodies mean size (Supp. Fig. 3B). phyB-GFP fluorescence rapidly localized into discrete nuclear foci after 2h exposure to light (Fig. 2G), with size of these foci gradually increasing throughout the day and their number reduced. Larger photobodies (>0.45-0.28 μm) declined after day-to-night transition (Supp Fig. 3D-F), and their disassembly into smaller foci was accelerated towards the second half of the night (Supp. Fig. 3G-I). At 28°C, the number of 0.45-0.28 μm photobodies per nuclei was severely reduced as compared to 22°C (Fig. 2G, Supp. Fig. 3D-F). This reduction, however, was not accompanied by a greater abundance of smaller foci during the day, but only during early night (Supp. Fig. 3G-I). Quantification of total nuclear fluorescence showed that phyB abundance is substantially reduced at 28°C (Fig. 3A), although diurnal changes in phyB-GFP fluorescence were not affected by temperature. Indeed, phyB-GFP levels remained roughly constant at 22°C in the day, and slowly declined at night, with an equivalent pattern observed at 28°C, except for a faster decline at night (Supp. Fig. 3A).

**Figure 3.**
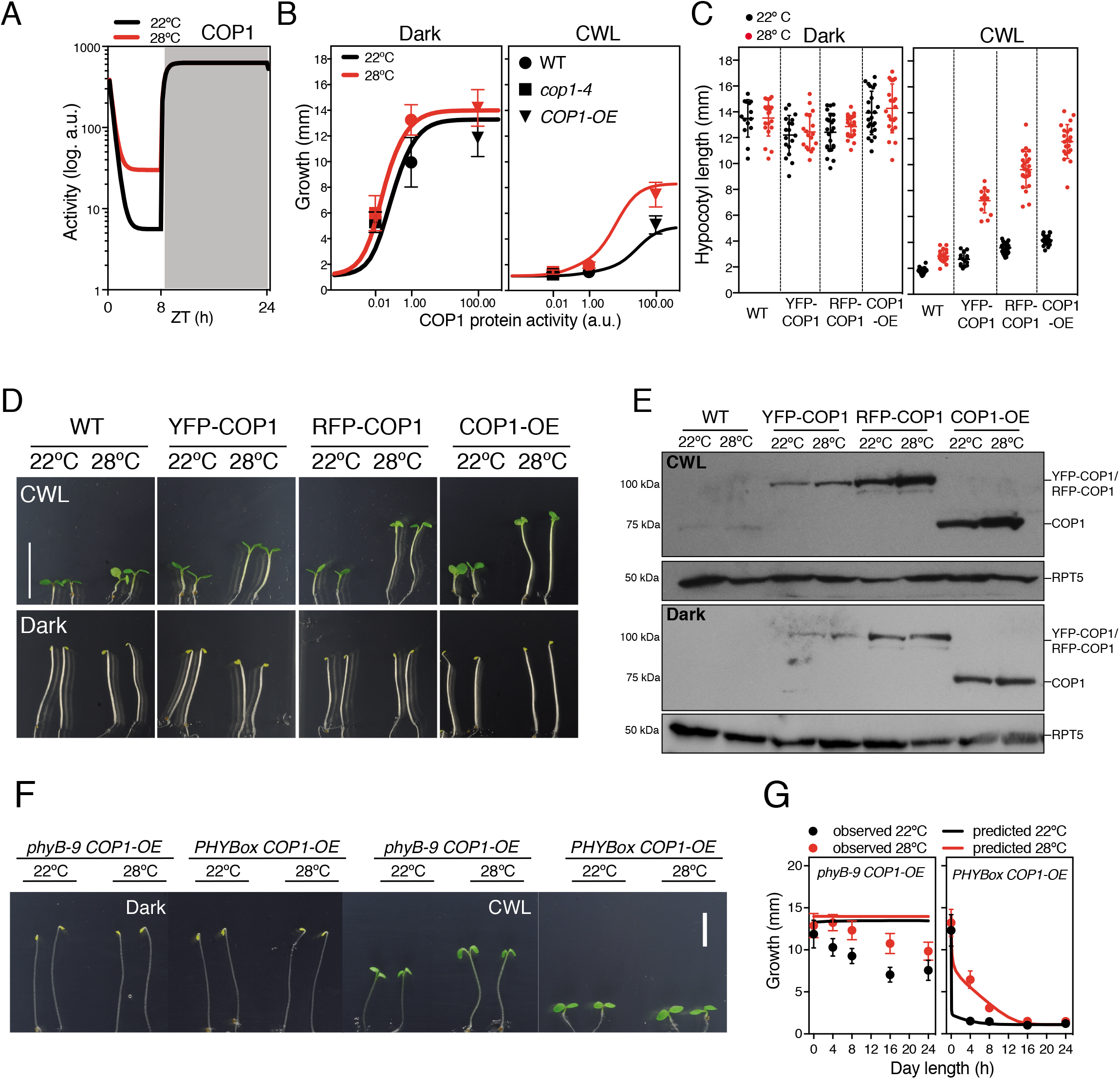
Effect of changes in COP1 activity on thermal growth. **(A)** Levels of COP1 activity predicted by the mathematical model in short day conditions. **(B)** Hypocotyl length as affected by the interaction of environmental conditions and COP1 levels. Observed hypocotyl lengths of Col-0, *cop1-4* and *COP1-OE* (symbols) versus the growth values estimated by the model (solid lines) in dark (left panel) and CWL (right panel). **(C)** Hypocotyl length phenotypes of different COP1 overexpressors. Bars indicate standard deviation (n=30). **(D)** Pictures of five-days-old seedlings grown in CWL (upper panel) and continuous darkness (lower panel) at 22°C or 28°C. Bar= 10 mm. **(E)** Protein blot showing that increased accumulation of the COP1 protein correlate with enhanced thermal elongation in CWL, but this response is saturated in darkness. COP1 was detected using an anti-COP1 antibody. RPT5 was used for loading control and detected using an anti-RPT5 antibody. Total protein extracts were used for immunoblot analysis. **(F)** Phenotypes of *phyB-9 COP1-OE* and *PHYBox COP1-OE* seedlings grown for five days either in darkness or CWL at 22°C or 28°C. **(G)** Observed hypocotyl growth (circles) of *phyB-9 COP1-OE* and *PHYBox COP1-OE*) lines grown at different day lengths and 22°C or 28°C. Solid lines are predictions from the model.

Temperature reduces phyB stability at a post-transcriptional level, since these lines expressed the phyB-GFP protein under control of the constitutive *35S* promoter. Related temperature-dependent changes in phyB stability were reported recently ^45^, although temperature treatments and results differed from ours. Active phyB levels steadily increased during the day in our growth conditions and reached maximal levels shortly before dusk, to exponentially decay at night (Fig. 2C). phyB was also found to be less abundant at 28°C, and large photobodies undergo fastest transition into smaller foci during early night (Fig. 2C). The model captures these dynamics, as it predicts that bioactive phyB is reduced at warm temperatures, and more rapidly inactivated in the dark (Fig. 2F).

### COP1 regulates thermomorphogenesis in the light

COP1 function is essential for thermal elongation ^31^, as reflected by the severely impaired thermoresponse of *cop1-4* genotypes (Fig. 1A). COP1 was shown to accumulate in the nucleus at 28°C, but higher activity in warm conditions to be uncoupled from light and timing information ^25^. Contrarily to these initial observations, our mathematical model predicts that COP1 activity is close to saturation and unresponsive to temperature at night, whereas its function is reduced during the day, being then critical for temperature-responsive HY5 turnover (Fig. 3A). We therefore simulated hypocotyl growth for a range of values of COP1 activity, and assigned the weak *cop1-4* alleles as a 100-fold reduction in activity, and *COP1-OE* lines as a 100-fold increase (Fig. 3B). Running the model with these arbitrary parameters captured thermal behavior of these genotypes, as it showed that reduced activity of *cop1-4* mutants leads to defective elongation, and an alike-impaired thermoresponse in darkness as in CWL. Ectopic COP1 expression, by contrast, is predicted to be of no effect in darkness, but lead to taller hypocotyls and an enhanced thermoresponse in CWL, as illustrated by a remarkable increase in hypocotyl length at 28°C (Fig. 3B).

Growth of *COP1-OE* and *hy5* lines, as compared to Col-0, follows this behavior (Fig. 1A; Supp. Fig. 1A and 1B), consistent with COP1 acting at temperature signaling in the light. We therefore analyzed the light-dependent thermoresponse of independent COP1 overexpressors by using *35S*::YFP-COP1 ^48^, *UBQ*::RFP-COP1 and *COP1-OE* ^49^ lines. Although these plants accumulate increasing levels of COP1 (Fig. 3C and 3D), their hypocotyl lengths were identical to the wild type in the dark, and failed to elongate further at 28°C (Fig. 3C and 3D). However, in CWL they displayed a thermal elongation response that was directly proportional to COP1 abundance (Fig. 3B). An identical behavior was observed in continuous red (CRL) and blue light (CBL) (Supp. Fig. 4A), in support of this response being independent of phyB Pfr thermal inactivation. Furthermore, COP1 abundance was reduced in etiolated seedlings (Fig. 3E), suggesting that COP1 auto-degradation is suppressed in the light.

Thermal Pfr inactivation is presumed to enable COP1-SPA1 re-assembly, and thereby favor COP1 activity in CRL. Modeling thermal behavior of *phyB-9 COP1-OE* and *PHYBox COP1-OE* lines actually predicted an identical growth phenotype for *PHYB*ox COP1-OE (Fig. 3F) and *PHYBox* plants (Fig. 1A), while *phyB-9 COP1-OE* lines to exhibit a constitutively tall phenotype, irrespective of temperature and day length conditions (Fig. 3G). Experimental validation of these phenotypes showed that *PHYBox COP1-OE* lines display, as predicted, short hypocotyls in the light and a hypersensitive response to day length thermal inhibition (Fig. 3G). However, they thermoelongate in short days and 4 h light cycles, indicating that temperature inactivates only a fraction of the Pfr pool and remaining Pfr exceeds COP1 and PIFs titration levels. On the other side, *phyB-9 COP1-OE* lines showed as predicted a tall phenotype in CWL, but retained a residual response to day length and temperature (Fig. 3F and 3G). These plants were in CWL taller than the rest of genotypes, but hypocotyl growth was partially suppressed as day length increased, and enhanced at 28°C. This behavior can be explained by the action of other photoreceptors not considered in the model. The blue light receptors cryptochromes (CRY1 and CRY2), for instance, disrupt COP1-SPA interaction, in addition to displace COP1 binding to its degradation targets via a conserved VP motif ^50–52^. CRY1 likewise interacts in a blue light-dependent manner with PIF4, suppressing PIF4-mediated hypocotyl elongation at elevated temperatures ^53^. In line with this notion, deviation of *phyB-9 COP1-OE* growth with respect to the model is accentuated in CBL, whereas significantly reduced in CRL (Supp. Fig. 4B). Opposing effects of red and blue light, thus support a phyB-independent role of COP1 in temperature signaling, this thermal role being additively enhanced by impaired phyB activity.

### Architecture of the thermomorphogenic network

Although the COP1 protein accumulates in the nucleus at elevated temperatures ^25^, biological relevance of this process remained an open question. Our modeling studies unveil a specific thermal signaling function of COP1 in the light, which most likely was masked in former studies. Fitting phyB thermal inactivation parameters ^4,5^ to the gathered growth data enabled us to derive a minimal thermal growth model where only the essential interactions were included. This model shows that the thermal responsive network comprises a triple feed-forward coherent motif, whereby ELF3 and phyB, on one hand, and COP1, on the other, exert opposing roles on thermomorphogenic growth (Fig. 1B).

Finally, we used this minimal model to explore the parameter space of these coherent motifs, to delimit their relative action and effects of day length duration on their thermosensory role. Heatmaps showing the contribution of COP1, ELF3, and phyB (horizontal axis) to thermal elongation (growth at 28°C compared to 22°C), and how these are predicted to vary with day length (vertical axis), are shown in Fig. 4. Also, linked effects of these motifs were inferred by generating similar heatmaps for the mutant backgrounds (as indicated in the panels). Red plot areas indicate conditions in which response to temperature is maximal, considering the combined effect of day length and activities of these regulators. For abscissa values=1, moving vertically on the ordinate axes shows that thermal elongation (color code on the right axis) is greater in short days (lower values in left axis), but significantly reduced in long days and CWL (upwards in left axis), as for our experimental data in Fig. 1A. Moving horizontally along a constant day length shows impact of changes in the different regulators on thermal elongation: i.e. for a 12 h day length, smaller abscissa values show that reduced COP1 levels abolish thermoelongation, while thermal growth is strongly enhanced in response to increased COP1 activity (higher abscissa values), in line with the *cop1-4* and *COP1-OE* phenotypes in Fig. 1A. An opposite behavior is observed for phyB activity, highlighting an opposing role of phyB and COP1 in driving thermal elongation. Thermal effects of high COP1 activity are lost in the absence of phyB, consistent with the phenotype of *phyB-9 COP1-OE* lines (Fig. 3F-G). Defective COP1 function moreover impairs thermoelongation under any condition (Fig. 4, middle row, central and right panels), highlighting the key contribution of this regulator to thermomorphogenesis.

**Figure 4.**
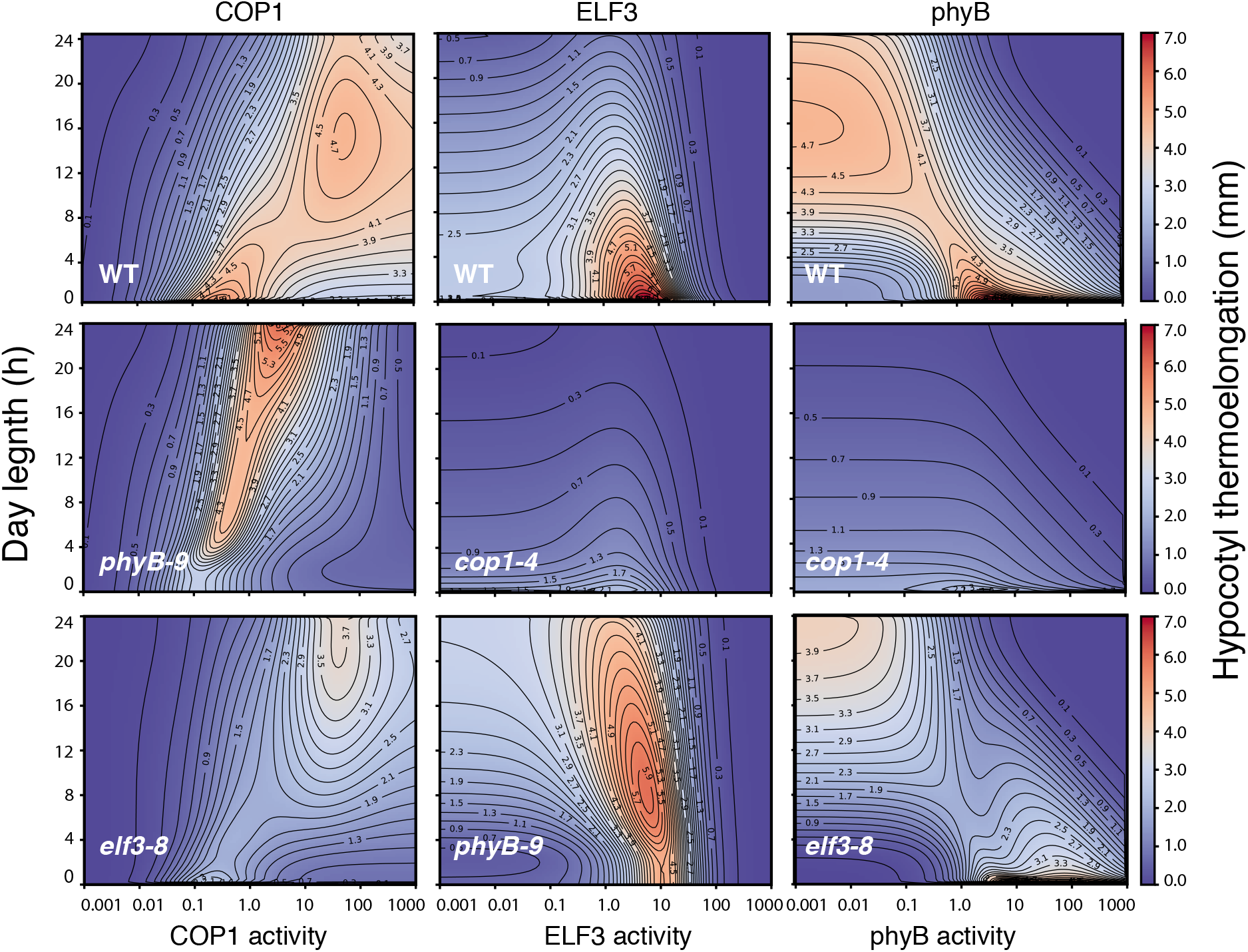
phyB, COP1 and ELF3 contribution to day length-dependent hypocotyl thermoelongation as predicted by the model. Heatmap plots representing hypocotyl thermoelongation (calculated as the difference of hypocotyl length at 28°C and 22°C) relative to day length (Y axis) and activity of the COP1, phyB and ELF3 proteins (X axis, as indicated). A value = 1.0 corresponds to wild type level, while greater and lesser values are equivalent to overexpression and loss-of-function, respectively. Backgrounds are indicated in the panels.

On the other side, ELF3 is predicted to have a main contribution to thermal growth in short days (Fig. 4, upper row, central panel), consistent with its role in driving *PIF4* expression during nighttime. Loss of ELF3 function, weakens thermal effects of elevated COP1 or reduced phyB levels and narrows them down to longer day lengths. Moreover, phyB is found to override ELF3 activity, as reflected by the enhanced thermal growth, over a wider range of day lengths, caused by elevated ELF3, in the *phyB-9* background (Fig. 4, lower central panel). Surprisingly, a small elevation in ELF3 is predicted to favor thermal elongation, whereas greater ELF3 levels reverse this positive effect (Fig. 4). LUX was actually shown to bind with high affinity the LBS recognition motif, whereas its activity would be abolished on dimerization with ELF3 and restored on additional interaction with ELF4 ^8^. It is thus conceivable that ELF3 counteracts LUX inhibition of *PIF4* and its own expression, until LUX elevation and higher EC levels reverse this positive effect. Even though this dual function remains to be substantiated, it is noteworthy that it is predicted to be potentiated by loss of phyB activity, hence underscoring a role of phyB signaling in antagonizing ELF3. Notably, COP1 thermal activity also appears in these simulations to be affected by the lack of ELF3, suggesting that ELF3 and COP1 function is somehow linked (Fig. 4, lower left panel). Thus, it will be interesting in future studies to validate these novel interactions and investigate their underlying molecular mechanisms.

## DISCUSSION

Thermal responsiveness is pivotal in nature to reduce the impact of potentially damaging temperatures. Warm-induced hypocotyl elongation drives the apical meristem above the warm soil, while upwards foliar orientation reduces the incidence of direct sun light and favors aeration to cool the leaves. Magnitude of this response is however reduced on exposure to full sunlight and longer day lengths of summer, which overlap with warmer temperatures. Understanding the molecular processes driving this inhibition is thus pivotal to optimize resilience to climate change of long day-requiring crops. In this work, we show that COP1 acts as a major thermal regulator in the light. COP1 was shown to accumulate in the nucleus at elevated temperatures ^25^, our findings pointing to warm temperatures interfere with nuclear exclusion of the protein in the light. The photoreceptors phyB, phyA and cryptochromes bind SPAs and disrupt COP1-SPAs complex formation ^19,54^. SPAs were reported to act as essential COP1 cofactors, being proposed to retain COP1 in the nucleus in the dark. This dual role has been however questioned by the observation that *spaQn* mutants normally accumulate COP1 in the nucleus in darkness, but fail at cytosolic COP1 relocation on exposure to light ^55^. Thereby, when unbound to COP1, SPAs may take part of the nuclear COP1 exclusion machinery, having a central role in COP1 thermal regulation. COP1/SPA activity suppresses light signaling by targeting degradation of several light response factors antagonizing PIFs, including HY5 and HFR1. The COP1/SPA complex targets also ELF3 ^26,28^ for degradation, and is required for enhanced *PIF4* expression and PIF4 protein stabilization in warm conditions ^25,56^. Therefore, it is conceivable that nuclear COP1 accumulation mediates enhanced thermal elongation through a combination of these processes.

As phyB and SPA1 interaction suppresses COP1 activity, it cannot be excluded that light-dependent COP1 thermal function is a primarily consequence of phyB Pfr thermal inactivation. Our results, however, do not support this mechanistic model. Loss of phyB or ELF3 function in the double *phyB-9 cop1-4* and *elf3-8 cop1-4* mutants is unable to rescue thermal elongation defects of *cop1-4*, revealing that COP1 activity is crucial for the enhanced thermal responsiveness of the *phyB-9* and *elf3-8* backgrounds (Fig. 1A). Positive effects of COP1 in CWL are additive to the *phyB-9* mutation, suggesting that thermal COP1 function is to an important extent independent of phyB activity. Yet *COP1-OE phyB-9* lines were not as elongated in CWL as predicted by the model (Fig. 3G), this divergence was notably reduced in CRL, while it was aggravated in CBL (Supp. Fig. 4B). This supports that this phenotype is caused by direct COP1 inhibition by the blue light receptor cryptochromes ^57,58^, calling for future improvement of the model, by incorporating these photoreceptors

Thermoelongation heatmaps (Fig. 4) predict that phyB and COP1 activities are more relevant as day length increases, while role of ELF3 is more prominent in short days. Both reduced phyB and higher COP1 levels lead to significantly enhanced thermal elongation in long days and CWL, while defective COP1 function abolishes thermal responsiveness irrespective of phyB and ELF3 levels. Therefore, the model fully captures the epistatic effects of *cop1-4* on loss of phyB or ELF3 function, supporting a pivotal role of COP1/HY5 in modulating PIF4 activity downstream of these regulators. Heatmap simulations also show that impaired ELF3 activity reduces phyB and COP1 thermal effects, consistent with defective EC function leading to a de-repressed response at 22°C. Model predictions thus substantiate the phenotypes of double overexpressor/ mutant lines, demonstrating that the modelled interactions are sufficient to explain *Arabidopsis* thermal behavior.

Temporal dynamics of ELF3 and PIF4 abundance, as measured using *pELF3::ELF3-LUC* and *pPIF4::PIF4-LUC* lines, showed that accumulation of these proteins, in particular PIF4, is strongly affected by day length. *PIF4* expression rises in short days during late night, while in long days it is elevated only during daytime (Fig. 2). Likewise, warm temperatures delay in long days the peak of *PIF4* transcription during late evening, leading to a marginal increase of PIF4 protein levels at dusk (Supp. Fig. 2I and 2H). COP1 was recently reported to destabilize the DELLA repressors at warmer temperatures ^59^ and in this regard, it is possible that this control takes on a central role in longer day lengths. While diurnal ELF3 oscillation was coarsely introduced when fitting the model, PIF4 activity was remarkably left unconstrained. However, in its current configuration the model correctly predicts that PIF4 activity is enhanced at elevated temperatures, while this effect is greater in short days as compared to longer day lengths. Thus, future up-grading of the model by using dynamics of these regulators as measured in the LUC lines, will certainly help to its fine-tuning and improve its already strong predictive power.

Most crop species show a narrow genetic variability in thermomorphogenic-related traits ^60^, which threats their productivity in a global warming scenario. Natural variation in the phyB and ELF3 loci was found to be associated with increased thermal sensitivity in *Arabidopsis* and several crops, in a consistent manner with their adaptation to local climate variables ^6,9,61,62^. Lower activity of these alleles results in an elongated phenotype that is also associated with earlier flowering, and thereby leads to a reduction in yield. We here show that COP1 activity is essential for increased thermal responsiveness of *phyB-9* and *elf3-8* mutants, and that COP1 over-expression leads to a significantly enhanced thermomorphogenic growth in the light, but a wild-type response in darkness. *COP1-OE* lines, on the other hand, do not exhibit the same accelerated flowering as *phyB-9* and *elf3-8* mutants. Hence, allelic variants in the *COP1* gene that lead to an overall increase in protein levels or activity emerge from this work as excellent candidates for optimizing crops resilience to climate change.

## MATERIALS AND METHODS

### Plant materials

*Arabidopsis thaliana* Col-0 plants were used as wild type (WT) in this study. The mutants and transgenic lines used in the different experiments were: *elf3-8* ^63^, *ELF3ox* ^26^, *pif4-101* ^64^, *pifq* ^65^, *phyB-9* ^66^, *COP1ox* ^26^ *COP1-OE* ^49^, *cop1-4* ^67^, *hy5-215* ^68^, *elf3-8 cop1-4* ^26^, *ELF3ox cop1-4* ^26^, *PIF4-HA* ^69^, *pELF3::ELF3-myc* ^7^, *pPIF4::PIF4-HA* ^42^, *cop1-4/* 35S:YFP-COP1 ^48^, 35S:PHYB-GFP ^70^.

### Growth conditions

Seeds were surface-sterilized for 15 minutes in 70% (v/v) ethanol and 0,1% (v/v) Tween 20, followed by two washes of 2 minutes in 96% (v/v) ethanol. Seeds were air dried and sown on half strength MS-agar plates with 1% sucrose and stratified for 3 days at 4°C in the dark. Germination was synchronized by illuminating the plates with white light for 4 hours and transferring them back to darkness for 20 additional hours. The seedlings were then grown in specific temperature and day length conditions with white light (50 μmolm ^−2^ s^−1^), red light (35 μmolm ^−2^ s^−1^) or blue light (35 μmolm ^−2^ s^−1^).

### Plasmid constructs and generation of *Arabidopsis* transgenic lines

To generate the p*ELF3*::LUC expression cassette, a fragment containing 2.21 kb upstream regulatory region of ELF3 was amplified from Col-0 genomic DNA using primers ELF3Prom-Fw (5’-CACCCTTATAAATAAAATTCC-3’) ELF3Prom-Rv (5’-CACTCACAATTCACAACCTTTTTC-3’) and cloned into the Gateway System (pENTR™ Directional TOPO^®^ Cloning kit, Invitrogen) to obtain the pELF3-TOPO plasmid. To generate *pELF3::ELF3-LUC* construct, the ELF3 coding region was amplified without the stop codon using the ELF3 cloning-Fw/ ELF3 cloning-Rv primers and cloned into pENTR™/D-TOPO vector. The PCR fragment of pELF3 sequence was amplified from pELF3-TOPO with ELF3 Prom NotI-Fw/ ELF3 Prom NotI-Rv and then inserted into *NotI* of ELF3-TOPO, to produce the *pELF3::ELF3-TOPO* plasmid. Finally, pELF3-TOPO and p*ELF3::ELF3*-TOPO constructs were mobilized by LR clonase II recombination (Invitrogen) into the pLUC-Trap3 vector ^71^.

To generate the p*PIF4*::LUC reporter gene, we amplified a fragment 2.47 kb upstream of the PIF4 ATG with primers PIF4Prom-Fw (5’-CACCCAGTACGCATCCAATCTTCTC-3’) and PIF4Prom-Rv (5’-CGGGATCCGGGTACAGACAGAAAGTGAC-3’) and followed the same procedure as described above for *pELF3::LUC*. The *pPIF4*::PIF4-TOPO construct ^42^ was transferred to the pLUC-Trap3 destination vector.

*COP1* CDS was amplified using primers COP1cloning-Fw (5’-CACCATGGAAGAGA TTTCGACGGATC-3’) and COP1cloning-Rv (5’-GAAGATCTTCACGCAGCGAGTACC AGAAC-3’) and cloned into the pENTR™/D-TOPO vector. The binary vector pUBN-RFP-Dest ^72^ was used for N-terminal fusion of the mRFP tag.

To generate *Arabidopsis* transgenic lines, the *Agrobacterium*-mediated floral dip method was used to transform p*ELF3*-LUC, p*ELF3*::*ELF3*-LUC, p*PIF4*-LUC and *pPIF4::PIF4-* LUC plasmids into Col-0, whereas pUBN-RFP-COP1 was transformed into *cop1-4* mutant background. Homozygous T3 plants were used in this study.

The double homozygous *phyB-9 COP1-OE and PHYBox COP1-OE* were generated by crossing the transgenic *COP1-OE* line with *phyB-9* and *PHYBox*, respectively. The phyB-9 mutation was genotyped using primers phyB-9-fw (5’-GCAA TGCCACACCTGTTCTTGTGG’3’) and phyB-9-rv (5’-CTTCACTAGGAGCAACACCCA ACG-3’). phyB and COP1 over-expression constructs were PCR-genotyped with primers 35S-fw (5’-CCACTGACGTAAGGGATGAC-3’) and phyB-9-rv and COP1-PE-fw (5’-CTTCCCTCCGTACTACACTCTTATC-3’), respectively.

### Hypocotyl measurements

Seeds were sown on half strength MS-agar plates with 1% sucrose and stratified 4 days at 4°C in darkness. After stratification, the germination was induced by placing the plates for 4 h under 50 μmol·m^-2^·s^-1^ white light and 20 h darkness at 22°C. After 4 days at the indicated photoperiod and temperature regimes, 20-30 seedlings were photographed and hypocotyl length was determined using ImageJ software.

### Luciferase activity assays

Luciferase activity assays were carried out using the LB 960 Microplate Luminometer (Berthold Technologies, UK). Transgenic *Arabidopsis* seedlings were plated on 0.5 x MS-agar medium and, after 2-3 days of germination, they were transferred to 96 well microplates (Costar, USA) filled with 175 μL 0.5x MS-agar medium 1% sucrose and 35μL (50 μg/mL) of Beetle Luciferin (Promega) dissolved in DMSO. Levels of luciferase activity were registered every hour and represented as the average of counts per two seconds in each well, with at least 24 plants per line. The luminometer was housed in a growth chamber to maintain the plants under controlled growth conditions.

### Confocal microscopy

Confocal fluorescence images of phyB-GFP were taken from the epidermis and the first sub-epidermal layers of the upper third portion of the hypocotyl. We used a LSM5 Pascal laser scanning microscope (Zeiss) with a water-immersion objective lens (C-Apochromat 40x/1,2; Zeiss). Probes were excited with an Argon laser (λ=488nm) and fluorescence was detected using a BP 505-530 filter. Images were taken from individual nuclei and image analysis was performed as in Legris *et al*., 2016. Each data point consists of the average of 6 replicates coming from two independent experiments showing the same trend. Replicates consist of the average of 3 plants coming from the same plate, and 3 nuclei per plant were analyzed.

### Western blot analysis

Total protein extracts from equal number of *Arabidopsis* seedlings were prepared by homogenizing plant material in extraction buffer containing 125 mM Tris HCl pH 7.4, 2% SDS, 10% glycerol, 6M urea and 1% β-mercaptoethanol. The extracts were heated at 95°C for 3 min and then centrifuged at 4°C for 15 min to recover the supernatant. Protein samples were boiled for 5 min in TMx2 loading buffer, and 40 μL of the protein extracts run on 8% SDS-PAGE gels. Homogeneous protein transfer to nitrocellulose membranes (Whatman) was confirmed by Ponceau red staining. For PIF4 and ELF3 detection, blots were respectively immunodetected with an anti-HA Peroxidase (Roche) or anti-myc antibody (Abmart), followed by incubation with anti-mouse HRP-conjugated antibody. Anti-RPT5 (Enzo) was used for equal loading control. For COP1 immunodetection it was used an anti-COP1 antibody kindly supplied by Dr. Hoecker ^73^ followed by incubation with a HRP-conjugated anti-rabbit secondary antibody. Chemiluminescence was detected with the Supersignal West Pico and Femto substrates (Pierce).

## SUPPLEMENTARY FIGURE LEGENDS

**Supplementary figure 1. Model predictions for hypocotyl thermoelongation of backgrounds or day length conditions different to those used to fit the model.**

**(A)** Growth of three *Arabidopsis* genotypes at either 22°C or 28°C under five white light day length conditions. **(B)** Growth of six *Arabidopsis* genotypes in a 4 h light regime and either 22°C or 28°C. Measured hypocotyl length (circles) were compared with values predicted by the thermoresponse model (solid lines). Lines are model predictions and not fits to the data. Bars indicate standard deviation (n=30).

**Supplementary figure 2**. **Effect of warm temperature on ELF3 and PIF4 expression and protein abundance under short and long days.**

**(A)** Total protein extracts from p*ELF3::*ELF3-myc transgenic lines were analyzed by Western blot using an anti-myc antibody. RPT5 was used for loading control and was detected using an anti-RPT5 antibody. Seedlings were grown for 7 days at 22°C or 28°C under short day conditions. Samples were harvested at the indicated ZT. Bioluminescence assays of Col-0 lines expressing ELF3-LUC **(B)** and PIF4-LUC **(G)** in short days. Bioluminescence recorded from transgenic seedlings expressing the p*ELF3::*ELF3-LUC **(C)**, p*ELF3::*LUC **(D)**, p*PIF4::*PIF4-LUC **(H)** and p*PIF4::*LUC **(I)** constructs in long days at 22°C/28°C. Values represent mean ± SE of absolute bioluminescence of at least 24 seedlings. (**E)** ELF3 and **(J)** PIF4 activity predicted by the model. **(F)** Western blot of p*PIF4::*PIF4-HA transgenic lines. The PIF4-HA protein was detected using an anti-HA antibody. Ponceau staining was used as loading control. Rectangles indicate light conditions: white, lights on, and grey, lights off. ZT indicates the zeitgeber time.

**Supplementary figure 3. phyB nuclear bodies formation is affected by temperature in short day conditions.**

*35S::PHYB-GFP* transgenic seedlings were grown under 50 μmol·m ^-2^·s^-1^ white light in short day cycles at 22°C or 28°C for 7 days. **(A)** Total nuclear fluorescence expressed in arbitrary units (a.u.). **(B)** Nuclear photobodies mean size (μm^2^). **(C)** Number of total bodies per nucleus. **(D-I)** Number of phyB nuclear bodies, sorted by size categories, in μm, measured with Matlab Software. The rectangles indicate the light conditions: white, lights on and grey, lights off. ZT, zeitgeber time.

**Supplementary figure 4. Thermal response of *COP1* lines in red and blue light.**

**(A)** Hypocotyl length phenotypes of the different COP1 overexpressor lines in continuous red (CRL) and continuous blue (CBL) light. **(B)** Phenotypes of *phyB-9 COP1-OE* and *PHYBox COP1-OE* seedlings grown for five days either in darkness, CWL, CRL and CBL, at 22°C or 28°C. Bars indicate standard deviation (n=30).

