## Supplemental Figures and Text for "Day length-dependent thermal COP1 dynamics integrate conflicting seasonal cues in the control of *Arabidopsis* elongation"

**Supplementary figure 1. Hypocotyl thermoelongation of mutant backgrounds.**

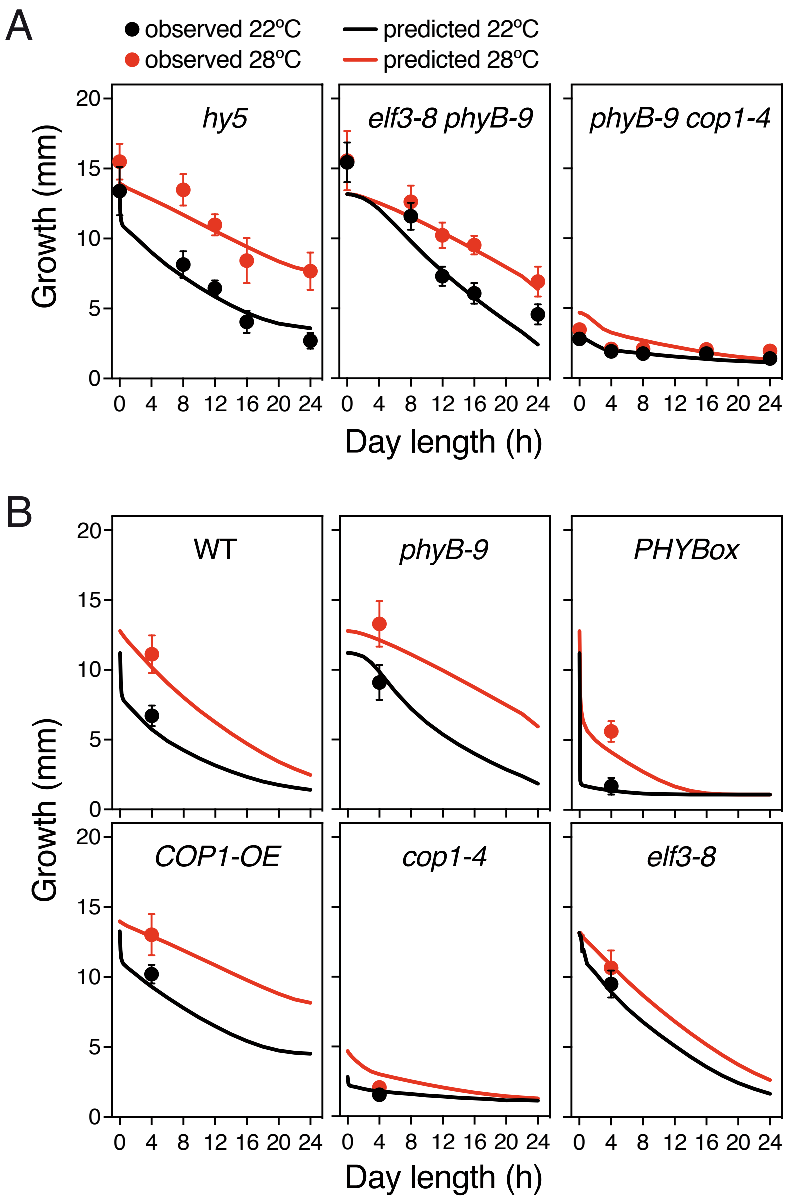


**Supplementary figure 1. Hypocotyl thermoelongation of mutant backgrounds.**
**(A)** Growth of three Arabidopsis genotypes at either 22ºC or 28ºC under five white light daylength conditions. **(B)** Growth of six *Arabidopsis* genotypes at 4 h light regime at either 22ºC or 28ºC. Observed measurements of hypocotyl length (circles) were compared with values predicted by the growth model (solid lines). Bars indicate standard deviation (n=30).

**Supplementary figure 2. Effect of warm temperature in ELF3 and PIF4 under short and long days.**

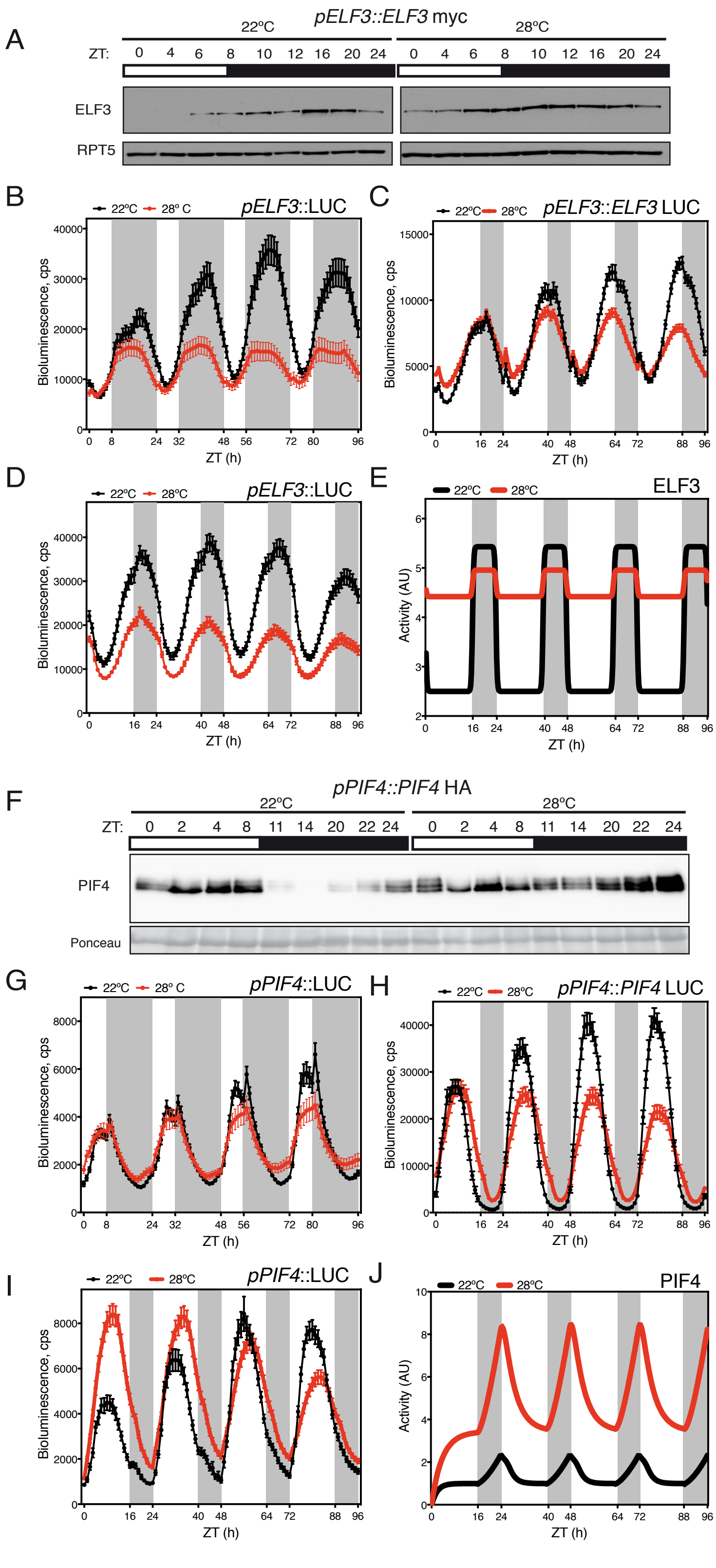

**Supplementary figure 2. Effect of warm temperature in ELF3 and PIF4 under short and long days.** **(A)** Total protein extracts from p*ELF3::ELF3-myc* transgenic seedlings were analyzed by Western blot using an anti-myc antibody. RPT5 was used for loading control and was detected using an anti-RPT5 antibody. Seedlings were grown for 7 days at 22ºC or 28ºC under short day conditions. Samples were harvested at the indicated ZT. Bioluminescence assays of Col-0 lines expressing p*ELF3-LUC* **(B)** and p*PIF4-LUC* **(G)** in short days. Bioluminescence recorded from transgenic seedlings expressing p*ELF3::ELF3-LUC* **(C)**, p*ELF3-LUC* **(D)**, p*PIF4::PIF4-LUC* **(H)** and p*PIF4-LUC* **(I)** constructs in long days at 22ºC/28ºC. **(E)** ELF3 and **(J)** PIF4 activity in long days predicted by the model. Values shown represent mean +/- SE of absolute bioluminescence of at least 24 seedlings. **(F)** Western blot of p*PIF4::PIF4-HA* transgenic line. PIF4-HA protein was detected using an anti-HA antibody. Ponceau staining was used as loading control. The rectangles indicate the light conditions: white, lights on and black/grey, lights off. ZT, zeitgeber time.

**Supplementary figure 3. phyB nuclear bodies formation is affected by temperature in short day conditions.**

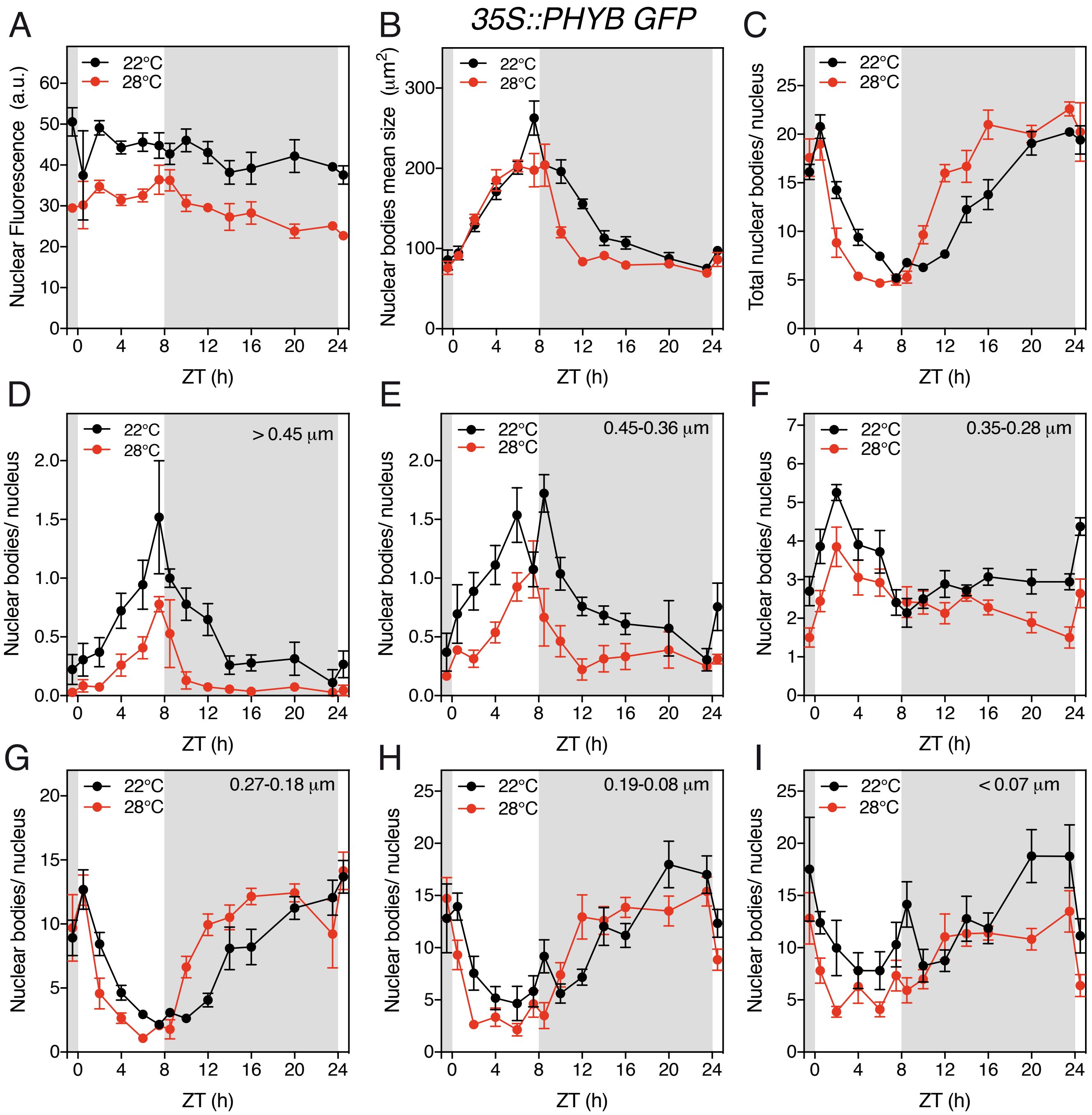


**Supplementary figure 3. phyB nuclear bodies formation is affected by temperature in short day conditions.**

35S::*PHYB-GFP* transgenic seedlings were grown under 50 µmol.m -2.s-1 white light in short day cycles at 22ºC or 28ºC for 7 days. **(A)** Total nuclear fluorescence expressed in arbitrary units (a.u.). **(B)** Nuclear photobodies mean size (µm2). **(C)** Number of total bodies per nucleus. **(D-I)** Number of phyB nuclear bodies, sorted by size categories, in µm, measured with Matlab Software. The rectangles indicate the light conditions: white, lights on and grey, lights off. ZT, zeitgeber time.

**Supplementary figure 4. Thermomorphogenesis in red and blue light.**

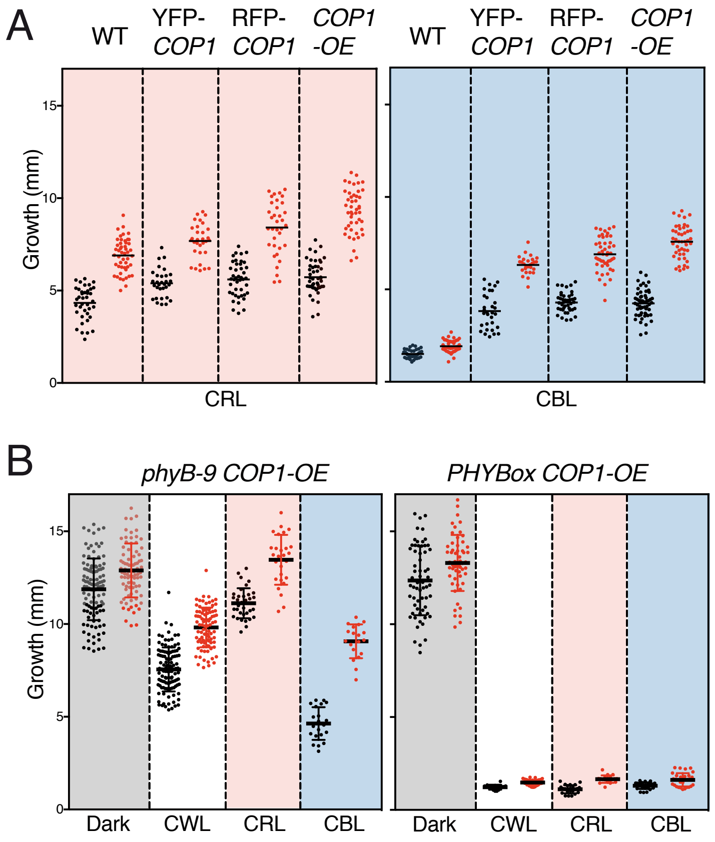


**Supplementary figure 4. Thermomorphogenesis in red and blue light.**
(A) Graphs showing the hypocotyl length phenotypes of different COP1 overexpressors grown in CRL and CBL. (B) Phenotypes of seedlings of *phyB-9 COP1-OE* and *PHYBox COP1-OE* grown for five days either in darkness, CWL, CRL and CBL at 22ºC or 28ºC. Bars indicate standard deviation (n=30).

### Day length-dependent thermal COP1 dynamics integrate conflicting seasonal cues in the control of Arabidopsis elongation

### Supplementary Text

Cristina Nieto, Pablo Catalán, Luis Miguel Luengo,
Martina Legris, Vadir López-Salmerón, Jean Michel Davière,
Jorge Casal, Saúl Ares and Salomé Prat

#### Code availability

All code and data to fit and simulate the mathematical model can be found at
https://github.com/pablocatalan/hypocotyl.

#### Mathematical model

##### Final model

The mathematical model used to obtain the results in the main text is as follows:

$$\begin{aligned}
\frac{dB(t)}{dt} &= p_B(T)L(t)(mut_B - B(t)) - k_r(T)B(t) \\
\frac{dE(t)}{dt} &= p_E(t,T,D,mut_E) - d_E E(t) \\
\frac{dC(t)}{dt} &= mut_C\left[p_{CL}(T)L(t) + p_{CD}(1-L(t))\right] - d_C C(t) \\
\frac{dP(t)}{dt} &= mut_P \frac{p_P}{1+p_{PE}(T)E(t)} - \frac{d_P}{1+k_{PC}C(t)}P(t) - d_{PB}B(t)P(t) \\
\frac{dG(t)}{dt} &= p_G + k_G \frac{p_{GP}P(t)}{1+p_{GP}P(t)+p_{GE}E(t)+p_{GB}B(t)+\frac{p_{GH}}{1+p_{HC}C(t)}},
\end{aligned} \tag{1}$$

where $t$ is time, $B(t)$ represents the concentration of the active form of phyB, Pfr, in the nucleus; $E(t)$ represents the concentration of ELF3 protein in the nucleus; $P(t)$ represents the nuclear concentration of PIF1, PIF3, PIF4 and PIF5; $C(t)$ represents COP1 in the nucleus, and $G(t)$ measures hypocotyl growth in mm, using the expression of PIF-targeted, growth related genes (e.g. *PIL1, XTH7* or *ATHB2*) as proxy. Parameters that are a function of $T$ (temperature) can have different values at 22°C and 28°C. $L(t)$ represents light, and it is either $0$ at night, or $1$ during daytime.

Parameters have the following meaning: for a given species $K$, $d_K$ is the decay rate of molecule $K$, $p_B$ is phyB's rate of activation and traslocation to the nucleus under light; $k_r$ is the rate of dark reversion, which following earlier reports [1] we assume happens during the day and night; $p_P$ is PIFs' production rate; $p_{PE}$ is the intensity of ELF3's inhibition of *PIFs* expression; $k_{PC}$ is the intensity of COP1's inhibition of PIFs degradation; $d_{PB}$ is the intensity of phyB's promotion of PIFs degradation and inactivation; $p_{CL}$ and $p_{CD}$ are, respectively, COP1's production rates during the day and the night; $p_G$ is the basal rate of hypocotyl growth; $k_G$ is the conversion between PIFs targets and growth; $p_{GK}$ is molecule $K$'s intensity of its effect on growth, $p_{GH}$ is related to the levels of HY5 (see below), and $p_{HC}$ is the intensity of COP1's inhibtion of HY5. Finally, $mut_K$ is a multiplier that alters molecule $K$'s production to acommodate knock-out and over-expressor lines; i.e. $mut_K = 1$ for the wild-type, $mut_K < 1$ for weak mutants and $mut_K > 1$ for over-expressor lines. In phyB and ELF3's case, knock-out mutants $phyB$ and $elf3-8$ have $B = 0$ and $E = 0$ at all times, respectively.

ELF3's expression follows a quasi-square wave:

$$p_E(t,T) = \begin{cases} mut_E p_{E1}(T) + p_{E2}(T) & \text{if } D = 0 \text{ hours.} \\ mut_E p_{E1}(T) - p_{E2}(T) \left(-1 + \frac{2}{1+\exp(-k_0 t_0)} - \frac{2}{1+\exp(-k_0 t_1)} + \frac{2}{1+\exp(-k_0 t_2)}\right) & \text{if } 0 < D < 24 \text{ hours.} \\ mut_E p_{E1}(T) - p_{E2}(T) & \text{if } D = 24 \text{ hours.} \end{cases} \tag{2}$$

where $D$ is the number of hours of light in the day; $p_{E1} + p_{E2}$ and $p_{E1} - p_{E2}$ are ELF3's average production in darkness and light, respectively; $t_0 = t \mod 24$, $t_1 = t_0 - D$ and $t_2 = t_0 - 24$, and $k_0 = 5$ h$^{-1}$. With this function, ELF3 oscillates between $p_{E1} + p_{E2}$ and $p_{E1} - p_{E2}$ rapidly. The advantage over using a simpler square wave is that this function is smooth, which prevents numerical anomalies. The value $k_0 = 5$ h$^{-1}$ defines the timescale of the rise and fall of the function when changing light conditions. It has been assigned arbitrarily to produce a smooth function but maintaining a sharp distinction between expression during light and darkness. Our results do not depend on the exact value, within reason, of this parameter.

We also assume that the over-expressor line $ELF3ox$ increases ELF3's production level $p_{E1}$, but that day-night oscillations, represented by $p_{E2}$ are not affected by the over-expression.

Parameter values can be found in Supplementary Table 1.

#### Development of the model

We have used the following experimental observations in order to develop the initial model:

1. phyB is activated by light and tends to spontaneously revert back to its inactive form. This 'dark reversion' is faster with higher temperatures. We follow the modeling in Jung et al. [2] of phyB's activation and dark reversion with small modifications.
2. phyB is marked for degradation by COP1, and this degradation constant is enhanced by PIFs [3].
3. phyB mediates the degradation of PIF4 and PIFs enhance the degradation of phyB [4, 5].
4. ELF3 is transcribed less during the day, and more during the night, in a sinusoidal pattern [6].
5. phyB physically interacts with ELF3, and this could potentially stabilize ELF3 [7].
6. COP1 also marks ELF3 for degradation [8].
7. PIF4 expression is inhibited by ELF3 as part of the evening complex [6]. This regulation is weaker at warmer temperatures, as the EC is impaired by temperature [9].
8. COP1 stabilizes PIF4 and PIF5 [10, 11].
9. COP1 is inactivated by phyB [12].
10. Hypocotyl growth is enhanced by the PIFs [13].
11. phyB prevents PIFs from binding to their targets [14].
12. ELF3 (independently from the evening complex) also prevents PIFs from binding to their targets [7].
13. HY5 represses hypocotyl growth, while COP1 mediates degradation of HY5 [15].

From these interactions, we developed the following model:

$$\begin{aligned}
\frac{dB(t)}{dt} &= p_B(T)L(t)(mut_B - B(t)) - k_r(T)B(t) - d_{BC}C(t)B(t) - d_{BP}P(t)B(t)\\
\frac{dE(t)}{dt} &= p_E(t,T,D,mut_E) - d_{EC}C(t)E(t) - \frac{d_E}{1+k_{EB}B(t)}E(t)\\
\frac{dC(t)}{dt} &= mut_C\left[p_{CL}(T)L(t) + p_{CD}(T)(1-L(t))\right] - d_C C(t) - d_{CB}B(t)C(t)\\
\frac{dP(t)}{dt} &= mut_P\frac{p_P}{1+p_{PE}(T)E(t)} - \frac{d_P}{1+k_{PC}C(t)}P(t) - d_{PB}B(t)P(t)\\
\frac{dG(t)}{dt} &= p_G + k_G\frac{p_{GP}P(t)}{1+p_{GP}P(t)+p_{GE}E(t)+p_{GB}B(t)+p_{GH}H(t)}.
\end{aligned} \tag{3}$$

Here $H(t)$ is HY5 concentration. As we do not have an equation for HY5, we assume it is in equilibrium and that its average levels are determined only by COP1: $H(t) = p_H/(1+p_{HC}C(t))$. The parameter $p_H$ is therefore no longer necessary and the effect of varying it is equivalent to variation of $p_{GH}$.

#### Simulated annealing

We wrote a custom simulated annealing algorithm [16] to fit Eqs. (3) to our experimental data (Fig. 1A, main text). We simulated the model for 5 days under all experimental conditions and for all mutants and tried to minimize an energy function that was the sum of all the squared errors between our experimental data (Supp. Fig. 5) and the model predictions. Together with growth data, we also used differences between model predictions and ELF3 levels in Col-0 and *phyB-9* backgrounds, Supp. Fig. 6. The energy function was

$$E = \sum_{k\in\mathcal{G}} (o_k - e_k)^2 + \sum_{k\in\mathcal{E}} (o_k - e_k)^2$$

where $\mathcal{G}$ and $\mathcal{E}$ are the sets of growth and expression datapoints, respectively. In other words, each datum contributed equally to the function. We used $3,544$ growth datapoints and 96 expression datapoints (see the Github repository for more details). We fixed $k_r(22) = 0.232$ h$^{-1}$ and $k_r(28) = 0.411$ h$^{-1}$, based on experimental measurements from Jorge Casal's lab, see Ref. [17]; $p_B(22) = 10$ in order to follow Ref. [2]; furthermore, $p_P$ and $p_{CL}(22)$ were set to 1 in order to reduce the dimensionality of the search space. All other parameters were set to 1 at the beginning of the search, allowing them to change ($p_{E2}(22)$ and $p_{E2}(28)$ were set to $0.9$ at the beginning of the search so as to allow variation in ELF3's production).

We started the process with all parameters set to 1. At each step $i$, we perturbed a randomly chosen parameter by adding to it a Gaussian random number with mean 0 and variance $0.1$, always checking that no parameter became negative. For this perturbed set of parameters, we computed the energy function $E_{\text{new}}$, compared it with the energy of the old parameter set $E_{\text{old}}$, and accepted the set with probability

$$p_{\text{acc}} = \begin{cases} 1 & \text{if } E_{\text{old}}/E_{\text{new}} \geq 1, \\ \frac{E_{\text{old}}}{E_{\text{new}}} T_A & \text{if } E_{\text{old}}/E_{\text{new}} < 1. \end{cases}$$

where $T_A = \dfrac{0.8}{\sqrt{1+i}}$; this particular form for $T_A$ was chosen after an initial trial-and-error stage. This means that the probability of accepting changes that increase the error decreased with each annealing step. A typical run for our model ran this process for $10,000$ steps, after which the variable $i$ was reset to zero and the process was started again, using as initial condition the final parameters of the previous run. This process was repeated $10$ times, to ensure the process did not get trapped in suboptimal minima. We then used the parameter configuration that minimized the energy function from among all visited configurations.

After obtaining a stable set of parameters for model (3), a few parameters in the model were close to $0$. New fits were made forcing these parameters to be zero, and the values obtained for the energy function were as good or even lower than those obtained considering the parameters free. The improvement in the fit when excluding these parameters can be explained by the increase in the efficiency of sampling the parameter space when its dimension is reduced. Finding this parameters consistent with zero in our fit does not mean that the interactions they represent do not exist, but rather that they are not important in our experimental conditions, or that their effect is already captured in an effective way by other parameters of the model. After exclusion of the parameters deemed negligible by our fitting procedure, the model given by Eqs. (3) is simplified to Eqs. (1). This final model was then fitted to the data using this simulated annealing procedure. Independent runs of this process (with the same initial conditions described above) converge to the same (or a very similar) set of parameters, shown in Supplementary Table 1.

#### Supplementary Figures

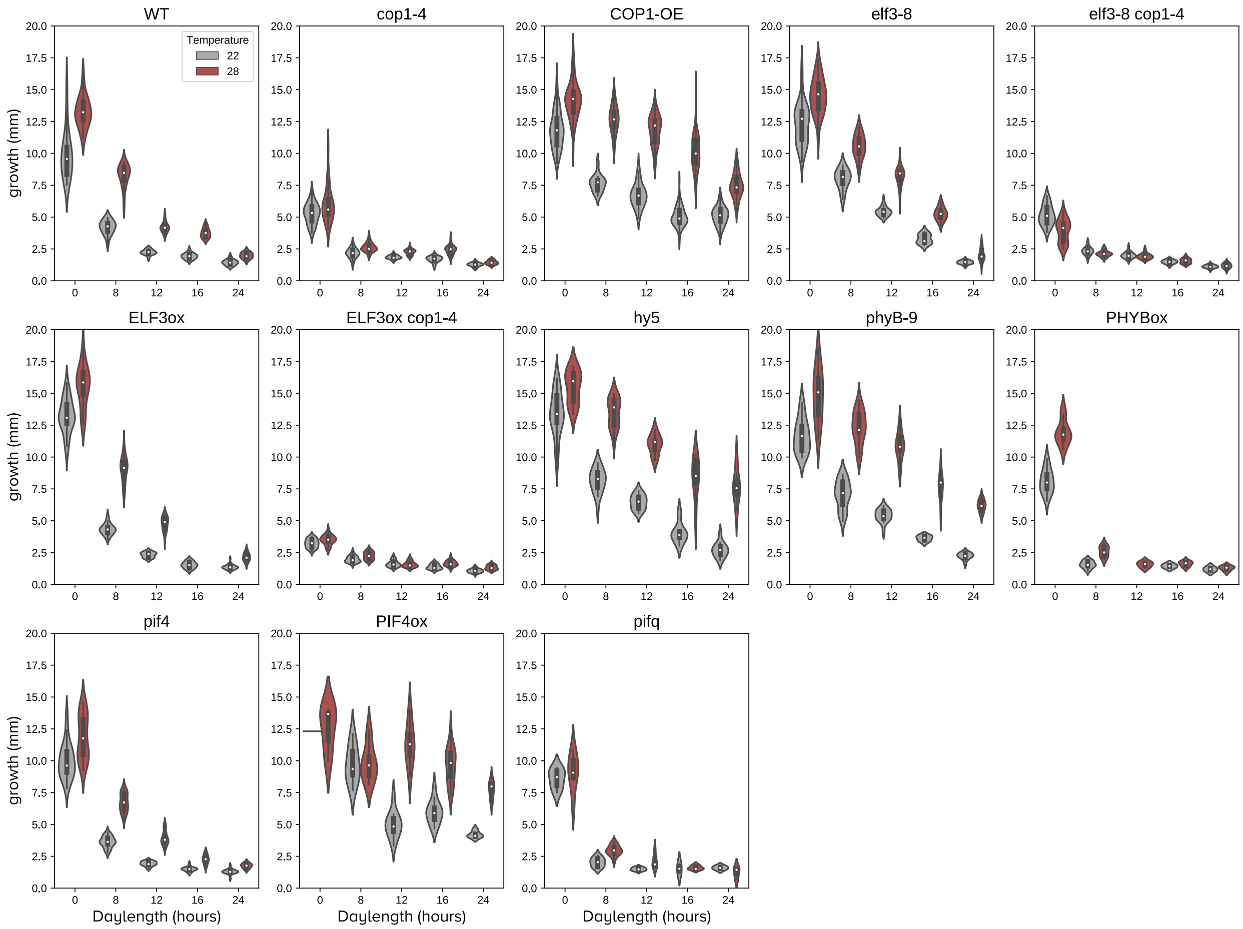


Supplementary Figure 5: **Experimental data used to fit the model (I).** We grew several mutant and over-expressor lines under different temperature and photoperiod conditions for five days (Methods), and used final hypocotyl length to fit our mathematical model.

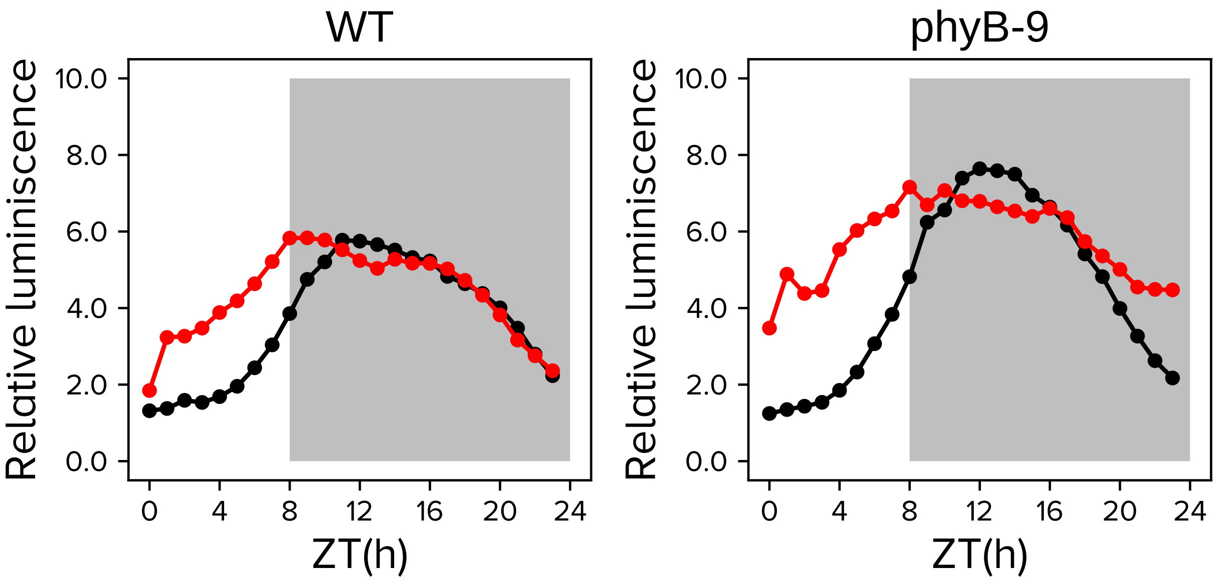


Supplementary Figure 6: **Experimental data used to fit the model (II).** We measured ELF3 levels using *pELF3*::ELF3 LUC transgenic lines under two different backgrounds: Col-0 and *phyB-9*, and used that information to fit our model.

#### Supplementary Tables

Supplementary Table 1: **Parameters used in Eq.** (1)**. The unit used for time is hours, concentrations are taken as non-dimensional.**

| | | | |
|---|---|---|---|
| $p_B(22)$ | 10.000 | $p_B(28)$ | 0.860 |
| $k_r(22)$ | 0.232 | $k_r(28)$ | 0.411 |
| $p_{E1}(22)$ | 107.721 | $p_{E1}(28)$ | 127.371 |
| $p_{E2}(22)$ | 39.828 | $p_{E2}(28)$ | 7.286 |
| $d_E$ | 27.172 | | |
| $p_{CL}(22)$ | 1.000 | $p_{CL}(28)$ | 5.370 |
| $p_{CD}$ | 112.374 | $d_C$ | 1.789 |
| $p_P$ | 1 | $d_{PB}$ | 0.313 |
| $p_{PE}(22)$ | 0.332 | $p_{PE}(28)$ | 0.028 |
| $d_P$ | 4.908 | $k_{PC}$ | 34.324 |
| $p_G$ | 0.009 | $k_G$ | 0.113 |
| $p_{GP}$ | 2.933 | $p_{GE}$ | 0.465 |
| $p_{GB}$ | 10.683 | $p_{GH}$ | 116.486 |
| $p_{HC}$ | 0.180 | | |
| $mut_B(phyB-9)$ | 0 | $mut_B(PHYBox)$ | 64.996 |
| $mut_B(elf3-8)$ | 0 | $mut_E(ELF3ox)$ | 1.185 |
| $mut_C(cop1-4)$ | 0.032 | $mut_C(COP1-OE)$ | 498.851 |
| $mut_P(pif4)$ | 0.495 | $mut_P(PIF4ox)$ | 6.367 |
| $mut_P(pifq)$ | 0.198 | | |
